# Construct validity of five sentiment analysis methods in the text of encounter notes of patients with critical illness

**DOI:** 10.1101/309195

**Authors:** Gary E. Weissman, Lyle H. Ungar, Michael O. Harhay, Katherine R. Courtright, Scott D. Halpern

## Abstract

Sentiment analysis may offer insights into patient outcomes through the subjective expressions made by clinicians in the text of encounter notes. We analyzed the predictive, concurrent, convergent, and content validity of five sentiment methods in a sample of 791,216 multidisciplinary clinical notes among 40,602 hospitalizations associated with an intensive care unit stay. None of these approaches improved early prediction of in-hospital mortality. However, positive sentiment measured by Pattern (OR 0.09, 95% CI 0.04 – 0.17), sentimentr (OR 0.37, 95% CI 0.25 – 0.63), and Opinion (OR 0.25, 95% CI 0.07 – 0.89) were inversely associated with death on the concurrent day after adjustment for demographic characteristics and illness severity. Median daily lexical coverage ranged from 5.2% to 20.5%. While sentiment between all methods was positively correlated, their agreement was weak. Sentiment analysis holds promise for clinical applications, but will require a novel domain-specific method applicable to clinical text.

## INTRODUCTION

In the era of widespread adoption of electronic health records (EHRs)[1] and learning health systems[2] there is growing interest in improving utilization of free-text data sources. Among patients with critical illness, the text of clinical notes has been used to identify diagnoses and interventions in the intensive care unit (ICU) and to improve predictions of future health states. [3–6] Clinical text contains important diagnostic information not found in structured data sources within the EHR.[7,8] But clinicians also make subjective assessments[9] and express attitudes about patient outcomes that may be purposefully or unwittingly inscribed in clinical notes. It is unknown if analysis of these subjective attitudes may augment existing yet imperfect mortality predictions,[10] improve communication by highlighting affective dynamics underlying patient-provider and patient-surrogate relationships, [11] or provide a feedback mechanism to clinicians regarding their implicit biases.[12]

The study of attitudes expressed in text is called “sentiment analysis” or “opinion mining.” [13] Dictionaries of terms (i.e. lexica) containing words with associated sentiment vary across different domains. [14] For example, “soft” may imply a different sentiment whether used with respect to sports or toys. [15] The analysis of sentiment in a medical context has been limited to patient opinions expressed in online social media[16,17] and in suicide notes, [18] the association of sentiment in hospital discharge documents with mortality and readmission, [19] and a descriptive comparison between nursing and radiology notes.[20]

Therefore, we sought to determine the construct validity of existing sentiment methods derived from other domains when used for analysis of clinical text among patients with critical illness. Specifically, we examined the predictive, concurrent, content, and convergent validity of these methods to assess different aspects of the sentiment construct.

## METHODS

### Population and data source

We analyzed the Medical Information Mart for Intensive Care (MIMIC) III database which comprises all hospital admissions requiring ICU care at the Beth Israel Deaconess Medical Center in Boston, MA, between 2001 and 2012.[21] Only hospital admissions with at least one clinical encounter note and a length of stay (LOS) ≤ 30 days were included.

### Text sources and sentiment methods

We aggregated clinical encounter notes at the patient-day level for each hospital admission and included notes from physicians, nurses, respiratory therapists, and other clinical specialties. We calculated the proportion of positive sentiment in each collection of daily aggregated notes as

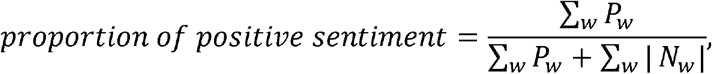

where *P_w_* and *N_w_* are the positive and negative sentiment scores, respectively, for each word iv in the daily aggregated text. We calculated separate scores using the Opinion,[22] AFINN,[23] EmoLex,[24] Pattern,[25] and sentimentr[26] methods. All five methods use simple dictionary lookups, and the latter two also account for valence shifters (e.g. “very” and “not”).

### Predictive validity

A sentiment measure with predictive validity should be strongly associated with some future outcome.[2 7] Therefore, for each method we trained a logistic regression model based on a random 75% sample of all hospital admissions to predict in-hospital mortality using data from the first day of the hospitalization. These were compared to a baseline model that did not include any sentiment measures. All performance measures were reported using the remaining 25% hold-out testing sample. The proportion of positive sentiment on the first hospital calendar day was the primary exposure, and the model was adjusted for age, gender, initial ICU type, modified Elixhauser score,[28,29] and initial sequential organ failure assessment (SOFA) score.[30] Model discrimination was assessed with the C-statistic and comparisons made with the DeLong test. [31] Calibration was assessed with the Brier score[32] and comparisons made using a bootstrapped t-test with 1,000 replicates.

### Concurrent validity

A sentiment measure with concurrent validity should be strongly associated with an outcome in the same time period.[27] Therefore, we examined the relationship between daily sentiment and the risk of mortality on the same day. We constructed a multivariable mixed-effects logistic regression model using the daily proportion of positive sentiment as the primary, time-varying exposure and daily risk of in-hospital death as the dichotomous outcome. The model was adjusted for age, gender, initial ICU type, and modified Elixhauser score.[28,29] A random effect was included for each hospital admission. A SOFA score (30) ≥ 7 was included as a dichotomous, time-varying exposure to account for daily changes in clinical severity. While daily SOFA scores have not been studied with respect to the daily risk of death, a time-varying score of ≥ 7 has been associated with an approximately 20% mortality rate in the ICU.[33]

### Convergent validity

The object toward which sentiment is directed (e.g. the patient, the prognosis, the tumor) may vary significantly. Each lexicon may also vary by the content of their terms and associated sentiment depending on the domain in which the method was developed and its original purpose. [20] Therefore, each sentiment method may provide a measure of some different aspect of the complex tapestry of sentiment found in clinical encounter notes. To assess the degree to which these five sentiment methods described the same phenomena, i.e. their convergence, [34] we measured their agreement with Cronbach’s alpha and calculated pair-wise Pearson correlations (*r*) at the patient-day level.

### Content validity

A useful construct of sentiment in clinical encounter notes should rely on keywords commonly used in the medical domain. Thus, the content validity is the extent to which a sentiment approach is capable of accounting for words and phrases found in these texts. [2 7] We measured this lexical coverage as the proportion of words in each patient-day’s aggregated text sample that was found in the lexicon.

Mixed-effects regression models were built using Stata (version 14.2, StataCorp, College Station, TX). Extraction of sentiment and training of other models were performed with the R language for statistical computing (version 3.3.2). The Pattern sentiment method was implemented using the Python programming language (version 2.7.13). We used a twosided alpha = 0.05 as a threshold for significance and adjusted all tests for multiple comparisons (Bonfeггoni correction). This study was considered exempt by the Institutional Review Board of the University of Pennsylvania.

## RESULTS

We analyzed 40,602 unique hospital admissions comprising 176,541 patient-days. The median hospital LOS was 3 days (Interquartile range [IQR] 2 – 5), the median age at admission was 60.8 years (IQR 39.4 – 75.8), and 3,728 (9.2%) patients died in the hospital. Each hospital admission contained a median of 8 (IQR 4 – 22) clinical encounter notes with median 1,480 words (IQR 588 – 5,289). These totaled 791,216 encounter notes containing 228,472,074 words. The distribution of daily sentiment for each method is summarized in Table 1.

The unadjusted temporal trajectories of sentiment stratified by in-hospital mortality are presented in Figure 1. However, the baseline model and all models with the addition of sentiment had C-statistic 0.79 without clinically relevant differences in discrimination (p = 0.06 – 0.76 for all comparisons). There were no meaningful differences in calibration with the addition of sentiment to a baseline model despite some comparisons achieving statistical significance (all models had Brier score 0.075; p = 0.005 – 0.481).

**Figure 1.**
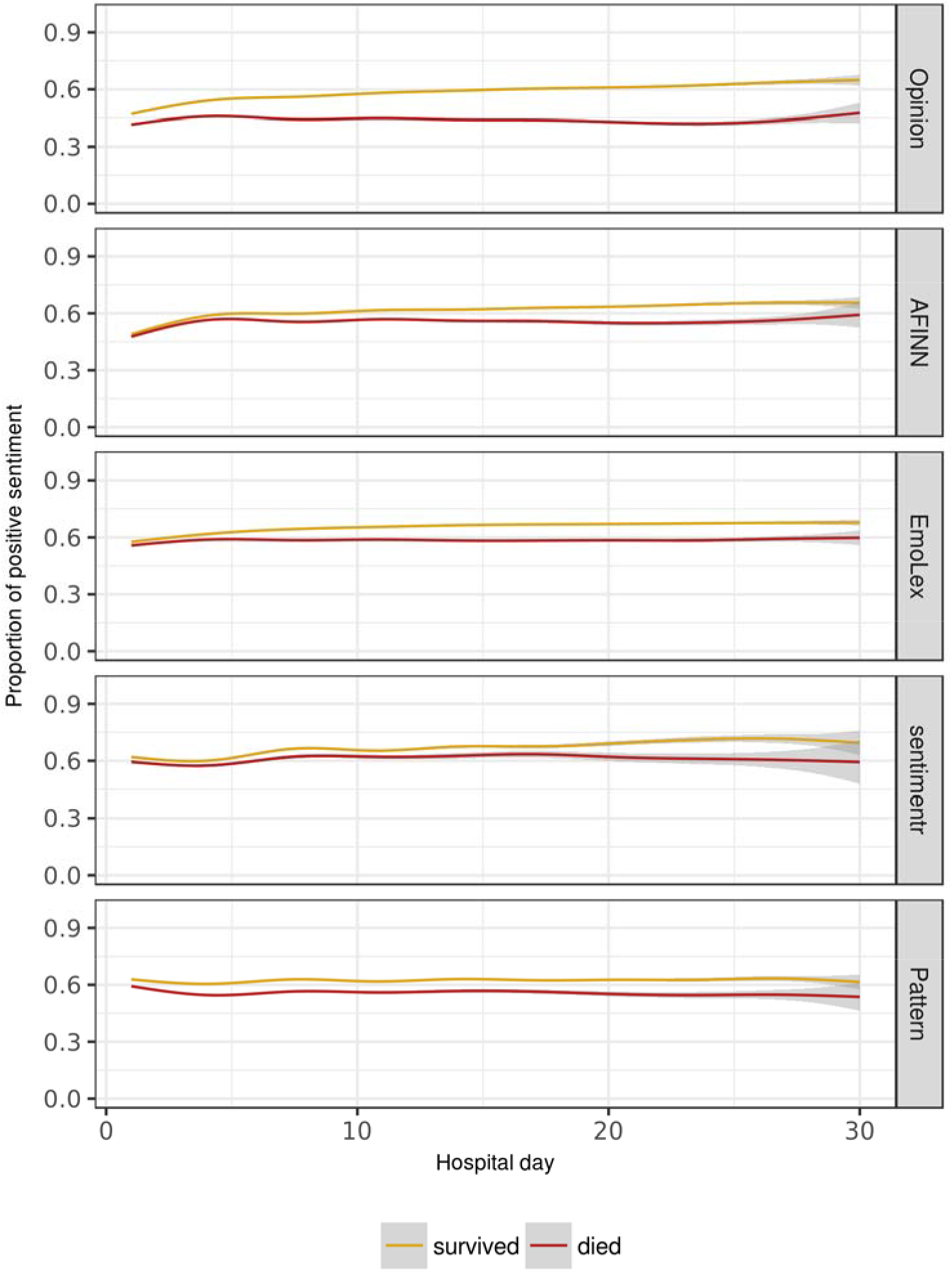
Unadjusted trajectories of sentiment in multidisciplinary clinical notes. Unadjusted trajectories of proportion of positive sentiment by sentiment method using a generalized additive model smoother with 95% confidence intervals

Sentiment was strongly associated with death when measured on the concurrent day for three of the five sentiment methods (Table 1). Even when adjusting for baseline characteristics and daily severity of illness, the proportion of positive sentiment measured by the Pattern method was inversely associated with the daily risk of death (OR 0.09, 95% CI 0.04–0.17).

As a measure of convergence, the Cronbach’s alpha for sentiment estimates for each patient-day was 0.65 (95% CI 0.64 – 0.65). All correlations between methods were positive and statistically significant, but most were of a modest magnitude (p<0.001; Figure 2). The median proportion of daily lexical coverage by hospital admission (Figure 3) ranged from 5.2% to 20.5%

**Figure 2.**
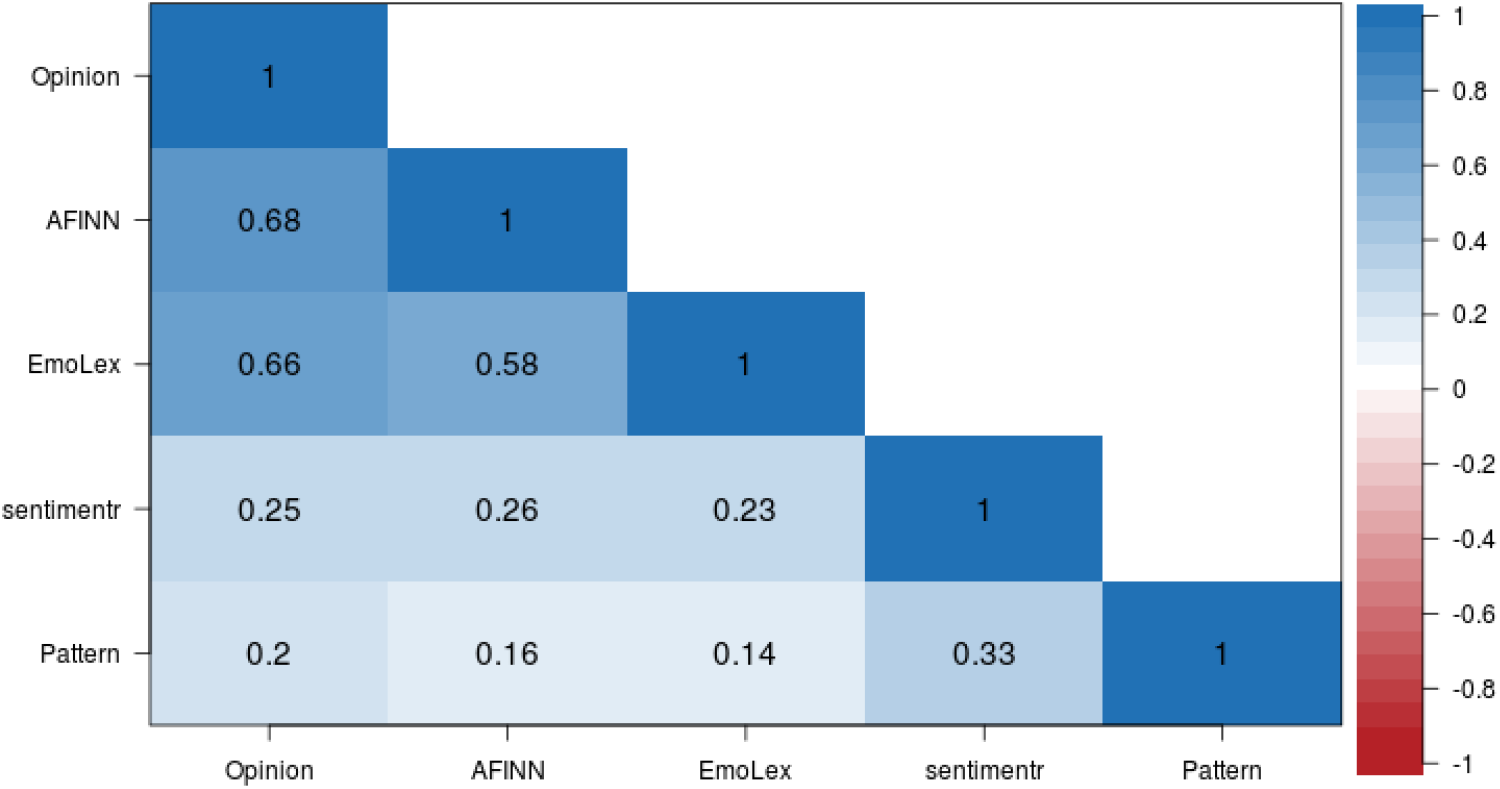
Pair-wise correlation between sentiment methods. Pair-wise Pearson correlations between methods of calculated sentiment by patient-day. AΠ estimates have p < 0.001 after adjustment for multiple comparisons.

**Figure 3.**
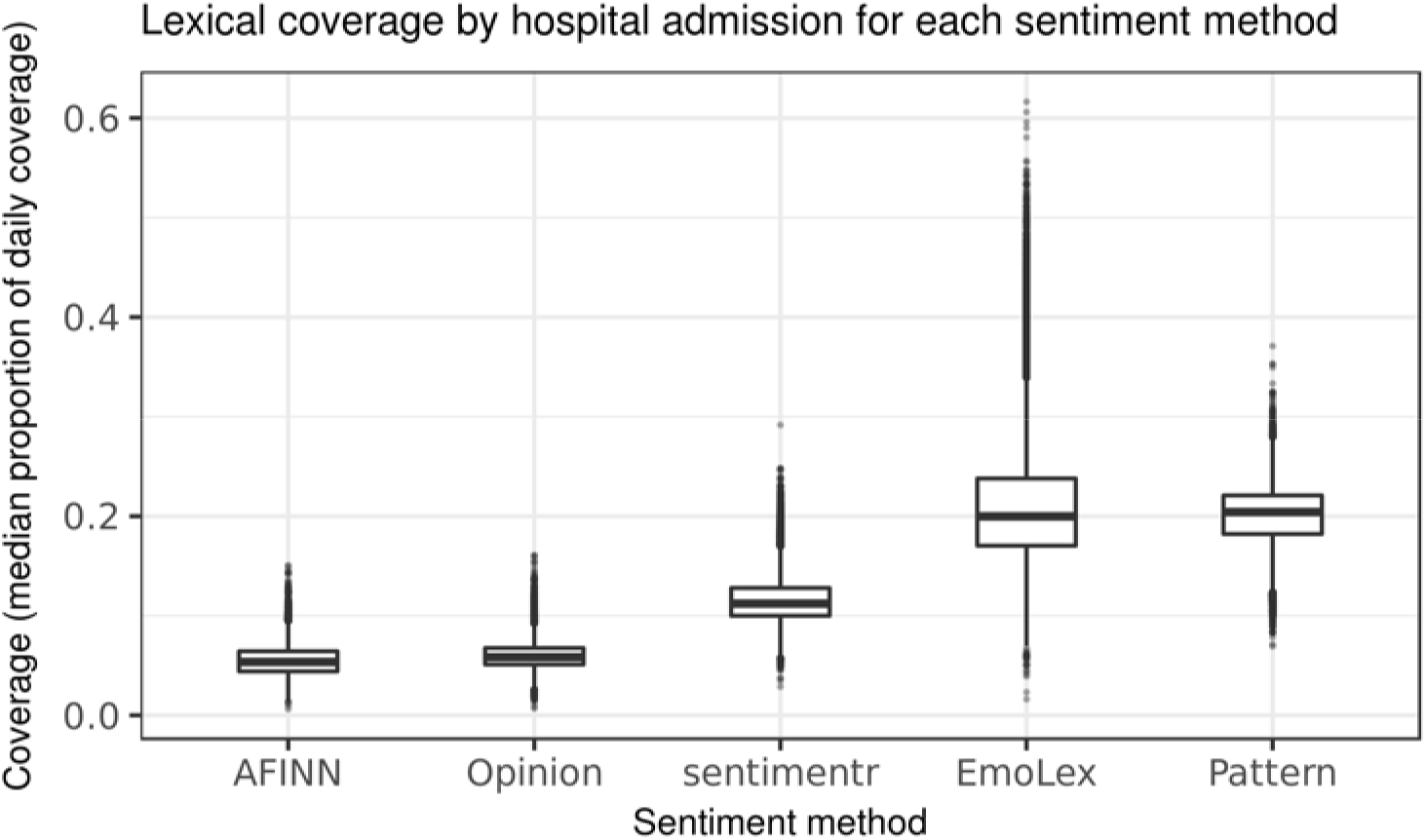
Lexical coverage by hospital admission for each sentiment method. The distribution of median proportion of covered words for each hospital admission by sentiment method.

The most common terms from the Opinion lexicon and representative samples of text are presented in Table 2. The associated polarity of these terms included instances with both concordant and discordant meanings in the medical domain.

**Table 1:** Adjusted odds ratio estimate for the proportion of daily positive sentiment for each sentiment method based on mixed-effects logistic regression model to assess concurrent validity; and distribution of daily sentiment.

| Sentiment method | Odds Ratio (95% CI) | p value | Median (IQR) |
| --- | --- | --- | --- |
| Opinion | 0.25 (0.07 - 0.89) | 0.033 | 0.50 (0.39 - 0.62) |
| EmoLex | 1.89 (0.41 - 8.69) | 0.412 | 0.60 (0.53 - 0.69) |
| AFINN | 0.65 (0.23 - 1.87) | 0.428 | 0.56 (0.44 - 0.68) |
| Pattern | 0.09 (0.04 - 0.17) | <0.001 | 0.63 (0.53 - 0.74) |
| sentimentr | 0.37 (0.25 - 0.63) | <0.001 | 0.71 (0.35 - 0.96) |

**Table 2:** The most common terms from the Opinion lexicon found in the clinical text sample and their polarity.

| Term | Polarity | Appearances (n) | Representative context |
| --- | --- | --- | --- |
| pain | negative | 658,808 | 'Pt reports back pain', 'Continue to monitor pain' |
| patient | positive | 588,213 | 'Encouraged patient to take his medicine', 'I saw and examined the patient' |
| stable | positive | 411,028 | 'stable frontal infarct', 'remains hemodynamically stable' |
| right | positive | 383,482 | 'only moving right arm', 'elevation of the right hemidiaphragm' |
| clear | positive | 368,261 | 'w/o clear evidence of infiltrates', 'Nutrition: clear liquids, advance diet' |
| well | positive | 365,899 | 'get radiation as well as this decision', 'satting well, no resp distress' |
| support | positive | 325,814 | 's/p arrest requiring ventilatory support', 'Emotional support given to patient & family' |
| soft | positive | 290,426 | 'abdomen soft slightly distended', 'possibility of soft tissue pus collection' |
| failure | negative | 268,838 | 'PNA with hypercarbic respiratory failure', 'R-sided heart failure leading to hepatopedal flow' |
| bs | negative | 259,638 | 'PULM: decreased bs on left', 'soft distended with hypoactive bs' |

## DISCUSSION

In our assessment of multidisciplinary encounter notes of patients hospitalized with critical illness, existing sentiment approaches demonstrated little evidence of most types of validity and exhibited high variability between methods. These results argue against the use of available sentiment methods to inform bedside clinical decisions, but also highlight opportunities to make sentiment methods more clinically applicable.

Many of the covered terms in this analysis had discordant polarity when applied in the medical domain. For example, the term “right” in medical parlance most often expresses anatomic laterality (e.g. “right ventricle”), thus should carry a neutral rather than positive sentiment with respect to prognosis. Similarly, the term “bs” is a shorthand abbreviation with multiple senses and may indicate “breath sounds”, “bowel sounds”, or “blood sugar” depending on the context. It should carry a neutral sense for all of these medical uses, but carried a negative polarity in the Opinion lexicon, where it may have been used originally to indicate a vulgar term in the online consumer reviews of electronics products.

The strong concurrent validity after adjustment for clinical and demographic characteristics suggests a temporal sensitivity of the sentiment to the patient’s clinical condition on the same the day. This finding was true even with adjustment for changes in severity of illness on each day, highlighting the presence of additional information encoded in free-text data not found in structured data sources such as laboratory values and vital signs. The models with the strongest effect sizes in this analysis (Pattern and sentimentr) were the only two that accounted for valence shifters. Nuances in expression of clinician sentiment are likely better captured by these approaches.

However, the addition of sentiment measures to a baseline prediction model resulted in no meaningful improvements to its discrimination or calibration. This lack of predictive validity suggests the temporal correlation of sentiment to mortality risk, while strongly concurrent, does not project forward in time.

Although all sentiment estimates were positively correlated with each other, their overall agreement was poor. The Opinion, AFINN, and EmoLex approaches were more highly correlated with each other (r=0.58 – 0.68), while the Pattern and sentimentr approaches were weakly correlated (r=0.33). These findings suggest a weak convergence towards two distinct constructs. More work is needed to distinguish between the sources, objects, and aspects of sentiment in clinical text. Additionally, the sentiment associated with objective medical terms (e.g. “cardiac arrest”) is distinct from the expression of a private state[35] of a clinician (e.g. “Mr. Jones is an unpleasant and uncooperative 65 year old man”). Each of these has separate analytic and ethical implications for use in clinical predictive modeling that have yet to be explored.

Finally, the content of sentiment lexica demonstrated coverage of medical terms that was higher than in previous analyses of medical text, but low compared to sentiment use in other domains. For example, Denecke et al. found coverage of 5% – 11% in radiology reports,6% – 11% in discharge summaries, and 8% – 12% in nursing notes, depending on the sentiment lexicon. [20] Coverage for the widely used SemEval Dataset range from 8% to 89%percent using commonly available sentiment lexica.[36]

The results of this study should be interpreted in the context of some limitations. First, the study analyzed data from a single academic center and may not generalize to the documentation style or patient population in other settings. Second, our analysis did not distinguish between the emotional valence of objective and subjective terms which conflates their practical use in clinical risk prediction.

In conclusion, this is the first study to examine sentiment in a set of multidisciplinary clinical encounter notes of critically ill patients and to assess the validity of these measures. Sentiment is strongly and concurrently associated with the risk of death even after adjustment for baseline characteristics and severity of illness. Our findings highlight the need for a domain-specific sentiment lexicon that has good coverage of medical terminology. Future work should seek to validate these findings in a broader population, better distinguish sources and objects of sentiment, and address potential ethical challenges of using sentiment to guide clinical care.

